# State or personality trait: Determinants of boldness and shoal size preference in adult zebrafish

**DOI:** 10.1101/2025.02.12.637681

**Authors:** Abhishek Singh, Prabhat Sharma, Mansha Jain, Pranati Mahajan, Jasmine Bajaj, Vaidehi Gupta, Niels J. Dingemanse, Bittu Kaveri Rajaraman

## Abstract

Animals adjust their behavior in response to physiological states to optimize the trade-off between energy acquisition and survival. A key question in animal personality research is whether behavioral variation is driven by consistent physiological state differences or stable personality traits. This study evaluates boldness and shoal size discrimination in adult male zebrafish (*Danio rerio*) across three hunger states: same-day feeding (H0), 24-hour food deprivation (H1), and 48-hour food deprivation (H2). Behavioral assessments were conducted over two testing cycles, with each individual undergoing every test twice for each hunger state within each cycle. The assays comprised the Open Field Test (OFT), Emergence Test (ET), and Social Preference Test (SPT), with the SPT conducted under binary (4 vs. 2 fish) and ternary (4 vs. 2 vs. 1 fish) choice conditions. Hunger significantly reduced emergence latency, but did not affect exploratory behavior in the OFT. Hunger also enhanced preference for larger shoals above chance in the binary SPT during the second testing cycle, though this effect was absent in ternary choices. Testing cycle effects revealed habituation-driven increases in boldness in both the tests and a progressive shift toward larger shoal preferences in the binary SPT. Among all assays, only the OFT showed significant repeatability, making it a reliable measure of boldness. However, it did not show consistent individual differences in plasticity due to hunger. These results highlight the crucial role of labile physiological states, such as hunger and habituation, in driving behavioral variation and underscore the importance of designing studies that disentangle state-dependent behaviors from stable personality traits in animal personality research.

## Introduction

Consistent differences in behavior between individuals within the same population, persisting across time and contexts, are termed personality, or temperament (Koolhaas et al. 1999; Réale et al. 2007; Dall and Griffith 2014; Laskowski and A. M. Bell 2014; Sih, Mathot, et al. 2015). These behavioral differences arise from genetic (Dingemanse, Both, et al. 2002; Van Oers et al. 2005; Sinn, Apiolaza, and Moltschaniwskyj 2006), epigenetic (Bossdorf, Richards, and Pigliucci 2008; Oers, Heuvel, and Sepers 2023), and neurodevelopmental (Bierbach, Laskowski, and Wolf 2017; Linneweber et al. 2020) mechanisms and have significant ecological and evolutionary consequences (Wolf and Weissing 2012; Jolles, Boogert, et al. 2017; Smith and Blumstein 2008). Commonly studied personality traits across species include boldness (Jolles, Briggs, et al. 2019), aggressiveness (Thys et al. 2017), sociability (Lucon-Xiccato and Dadda 2017), exploration (Schuett et al. 2013), and activity (Bailey et al. 2021; H. R. Thomson et al. 2020). Between individual correlations of these behaviors across contexts and time, known as behavioral syndromes (Sih, A. Bell, and Johnson 2004; Dingemanse, Kazem, et al. 2010; Jandt et al. 2014; Dingemanse, Dochtermann, and Nakagawa 2012), may result from hormonal (McGlothlin and Ketterson 2008) and genetic constraints, and can emerge through selection favoring specific combinations of traits (Dingemanse, Wright, et al. 2007; Adriaenssens and Johnsson 2013).

Individuals adjust their behavior in response to external conditions and fluctuating internal physiological states, exhibiting behavioral plasticity. This plasticity can be both repeatable and heritable (Mathot and Dingemanse 2014; Stamps 2016; O’Dea, Noble, and Nakagawa 2022; Araya-Ajoy and Dingemanse 2017; Mitchell and Biro 2017). State-behavior adaptive feedback loops, driven by interactions between physiological states (internal factors shaping optimal behavior) and resulting behaviors, have been proposed as key mechanisms underlying these consistent individual differences (Sih, Mathot, et al. 2015). Studies on individual behavioral consistency must account for an individual’s bodily state. Neglecting state-dependent effects can lead to erroneous conclusions: either falsely identifying a lack of consistency when states fluctuate significantly between trials, or mistakenly attributing consistency to personality when it is driven by state differences (Näslund and Johnsson 2016). Few studies have actively controlled for or manipulated internal states when measuring behavioral repeatability, and those that have often reveal only a weak or no relationship, making it challenging to differentiate between behavioral variation driven by stable personality traits and that influenced by state-dependent effects (Parthasarathy et al. 2022; Rádai et al. 2022; Näslund and Johnsson 2016; Moran et al. 2021).

Boldness is the most commonly studied personality trait, but also remains most ambiguously defined (Réale et al. 2007; Alecia J Carter et al. 2012; Conrad et al. 2011). Boldness is defined as an animal’s response to a risky situation (Réale et al. 2007) or the propensity to take risk (Coleman and Wilson 1998). Studies on boldness have been broadly categorized into measuring an animal’s response to a novel environment, a novel object, or predation risk (Christina N. Toms, David J. Echevarria, and Jouandot 2010). Due to the popularity of the test and its ambiguous definition, the assessment of boldness is susceptible to the jingle-jangle fallacy, where a similar behavior in slightly different versions of the test might be measuring two different traits (jingle) or a single trait may be measured under different labels (jangle) (Alecia J. Carter et al. 2013). Inferring personality based on a single behavioral test lacks test validity, emphasizing the need for multi-trait, multi-test experimental designs to determine whether the tests measure the same or different traits (Alecia J. Carter et al. 2013).

Apart from personality traits, there is growing interest in studying interindividual variation in cognitive abilities such as learning, behavioral inhibition, attention, and decision-making, as well as exploring whether these abilities correlate with personality traits (Dougherty and Guillette 2018; Griffin, Guillette, and Healy 2015; Guillette et al. 2015; Fu, Zhang, and Fan 2024). Shoal size preference tasks in fish, where individuals choose between two shoal size options, are often used to study numerical competence (Bossdorf, Richards, and Pigliucci 2008). These tasks are thought to involve complex cognitive processing compared to simpler behaviors like the tendency to shoal with conspecifics in a single-option task, known as sociability (Gartland et al. 2022). While the shoal size preference task with two shoal choices is well-established in fish behavior research (Agrillo, Dadda, and Bisazza 2007; Seguin and Gerlai 2017; Lucon-Xiccato et al. 2016), recent studies have expanded this approach to a three-shoal preference task to explore multi-alternative decision-making and the decoy effect (Reding and Cummings 2019; Singh, Kumari, and Rajaraman 2022). The decoy effect, defined as a shift in relative preference between two options upon the introduction of a third, irrelevant option, has been studied in various animal models (Latty and Trueblood 2020; Parrish, Evans, and Beran 2015; Orlando et al. 2023; Hemingway, DeVore, and Muth 2024; Bateson, Healy, and Hurly 2002; Cohen et al. 2019).

Zebrafish have become a prominent model organism for geneticists and neurobiologists, supported by a broad array of advanced experimental tools. These tools include molecular genetic techniques (Varshney et al. 2015; Klatt Shaw and Mokalled 2021), pharmacological methods (Agrillo and Bisazza 2014; Shams and T Gerlai 2016), and neurobiological approaches (Rinkwitz, Mourrain, and Becker 2011; Mrinalini et al. 2023). Measuring personality in zebrafish is of profound interest (Moretz, Martins, and Robison 2007; Oswald, Singer, and Robison 2013; Way et al. 2015; Christina N Toms and David J Echevarria 2014; Rey, Digka, and MacKenzie 2015; Cresci et al. 2018; Baker et al. 2018; Hamilton et al. 2021) as also is the relationship between personality and cognitive abilities (Lucon-Xiccato, Montalbano, and Bertolucci 2020; Daniel and Bhat 2020; Varma et al. 2020; Fu, Zhang, and Fan 2024; Corcoran, Storks, and Wong 2024) as well as the relationship between physiological state and personality (Araujo-Silva et al. 2018; Ariyomo and Watt 2015; Oswald, Drew, et al. 2012; Beigloo et al. 2024; Polverino et al. 2016). However most of these studies suffer from common malpractices of study design and use of statistics to measure personality like; the use of a single measurement of behavior to infer personality, using unpartitioned phenotypic associations to infer relationships with state or cognitive abilities and measuring phenotypic correlations without separating between-individual and within-individual variance components. A separate line of studies have shown the effect of hunger state on behavioral measures (Dametto et al. 2018), perception and decision making in zebrafish in both prey-predator (Chathooth et al. 2024; Filosa et al. 2016; Zaupa et al. 2024) and social context (Jens Krause, Hartmann, and Pritchard 1999). However, to our knowledge, no study has directly investigated the effects of hunger and habituation on shoal size preference in zebrafish, while also examining its repeatability and the potential behavioral syndrome between boldness, exploration, and shoal preference. In addition, we extend upon the shoal size preference assay with a the novel three option shoal size preference task, to study repeatability and effect of hunger state and habituation on multialternative decision making.

We conducted a fixed sequence of behavioral tests on the same set of fish: an Open Field Test (OFT), an Emergence Test (ET), a Social Preference Test (SPT) with two options (binary), and a three-option version (ternary). All tests were completed within a single day, and this battery was repeated twice, spaced two days apart, for each of three hunger states: same-day fed (H0), fed a day prior (H1), and fed two days prior (H2). Two such testing cycles were conducted, with each hunger state tested twice per cycle, to disentangle the effects of hunger from potential behavioral plasticity due to isolation. With this study design, we could ask the following questions.

1. Do individuals differ in their average performance in the open field test, the emergence test, and binary and ternary social preference test?
2. Do individuals differ in their plasticity to changes in hunger state or habituation due to testing cycles in each of these tests?
3. Is individual performance correlated between tests over multiple sampling events?
4. Does hunger state affect the mean behavioral performance in these tests?
5. Do trials repeated within each hunger state and cycles spaced by weeks affect the mean behavioral performance in these tests?

## Methods

### Subjects and Housing

Adult zebrafish (*Danio rerio*) of the same age group (3–6 months) were procured in two population cohorts from the local pet store in Daryaganj, New Delhi, India. The fish were housed for over a month before the experiment at Ashoka University, Sonipat, Haryana, India, in a ZebTec Active Blue - Stand Alone system (Tecniplast, PA, USA). They were maintained under a 12:12 light-dark cycle (10 a.m. to 10 p.m.) with water parameters set at a pH of 7.50–8.50, temperature of 28–30°C, and conductivity of 650–700 µS. The fish were fed powdered Tetra-Tetramin flakes ad libitum daily at 10:30 a.m.

Fish were moved to individual tanks (2 L volume) within the same fish system and maintained under identical conditions for one week prior to the experiments to enable individual identification. Due to time constraints allowing only a limited number of tests during daylight hours, the experiments were conducted in batches of six fish each. Four such batches were performed over four months to achieve a total sample size of 24. However, one fish died during the Batch 2 experiments, so an additional fish was included in Batch 3, bringing the number in that batch to seven. Only male fish were used in the experiments to control for sex-based differences in boldness (Mustafa, Roman, and Winberg 2019) and social preference (Velkey et al. 2022).

### Boldness Assays

We used two independent tests to measure boldness: the open field test (OFT) and the emergence test (ET). We avoided combining the two tasks involving both a shelter and an open arena for exploration, as this could capture the same inter-individual variation twice (e.g., individuals that emerge first are also likely to explore the arena more extensively), potentially leading to the artifact of a pseudo-syndrome.

### Open Field Test

The open field test was conducted in a glass tank measuring [45cmX18cmX30cm] with water filled till 15 cm height, equipped with a transparent removable plexiglass barrier to restrain the fish in the tank’s center (see apparatus, Figure 1). The tank was illuminated internally using LED strips affixed to its sides, while the walls were lined with black paper to minimize external light interference from the darkened laboratory setting. The arena was recorded using an overhead camera (Lenovo 300 FHD 1080P 2.1 megapixel webcam), securely mounted on a tripod at a fixed distance above the tank. A laminated grid sheet with 1 cm intervals was placed at the bottom of the tank to facilitate measurements.

**Figure 1.**
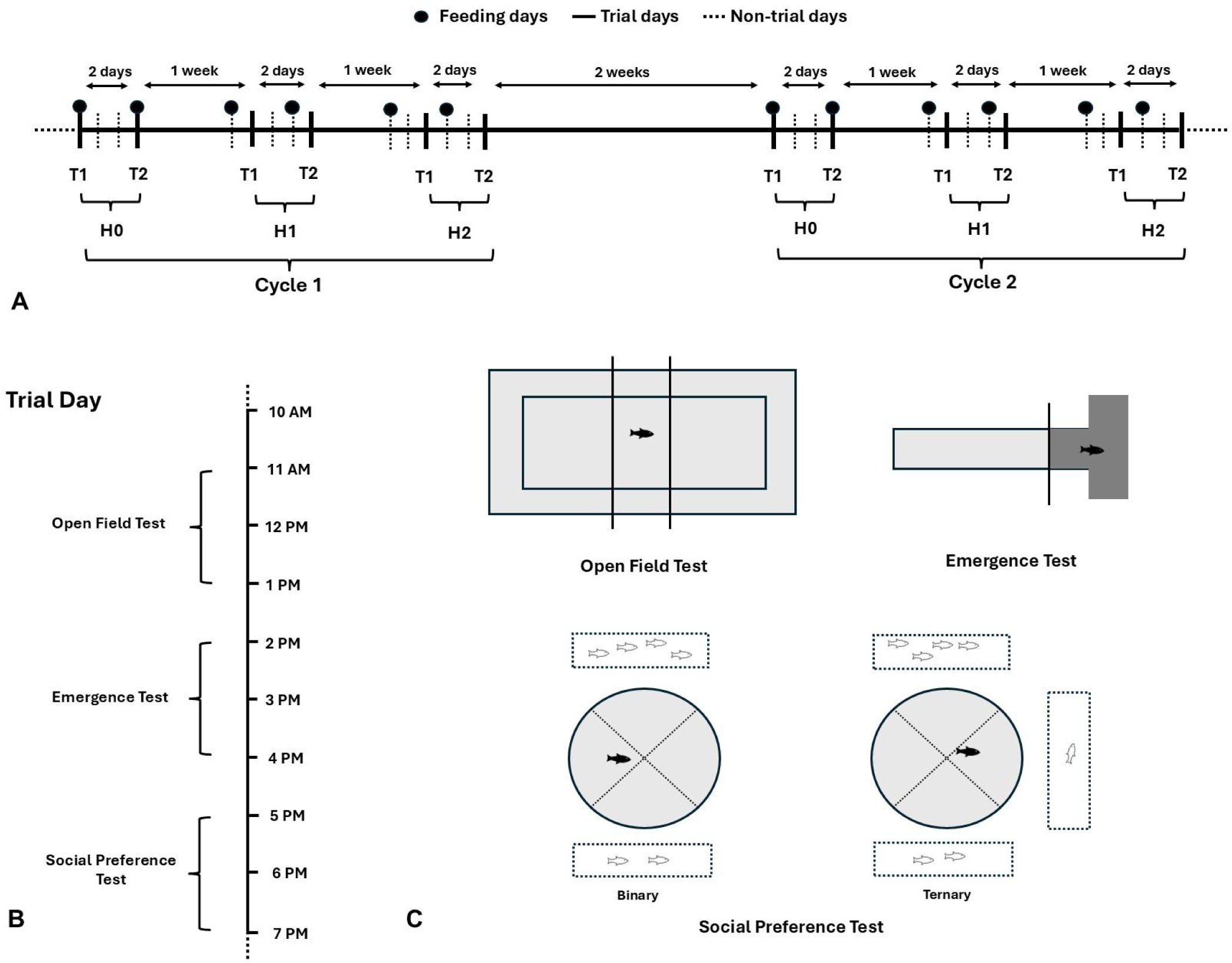
Experimental design, trial day timeline, and behavioral setups (A) Overview of the experimental design, showing two cycles (Cycle 1 and Cycle 2) consisting of trial (T1, T2) and non-trial days. Feeding days are marked with black circles. (B) Timeline of a typical trial day, demonstrating the sequence of behavioral tests conducted at specific time intervals: Open Field Test (10 AM–1 PM), Emergence Test (2 PM–4 PM), and Social Preference Test (5 PM–7 PM). (C) Schematic representation of the behavioral setups: Open Field Test, Emergence Test, and Social Preference Test in binary and ternary conditions.

During the open field test (OFT), fish were placed in a glass tank and confined to a central area for a 2-minute acclimation period. The barrier was then removed, allowing the fish to explore the open field arena for 3 minutes. Videos were analyzed using BORIS behavioral event logging software (Friard and Gamba 2016) to record state events of time spent in the open field. The open field was defined as the area beyond a 2 cm margin from the tank edges (Figure 1).

### Emergence Test

The emergence test (ET) was conducted using a modified T-maze (see apparatus, Figure 1). The maze consisted of two short side arms, each measuring 10 cm in length and 10 cm in width, connected perpendicularly to a longer central arm measuring 40 cm in length and 10 cm in width. A 10 cm section of all three arms at the junction was covered to create a dark holding zone, while the remaining 30 cm of the central arm was fully illuminated to serve as an exposed arena. The maze was filled with water to a height of 15 cm. A black guillotine door separated the dark holding zone from the open passage. The arena was recorded using an overhead camera (Lenovo 300 FHD 1080P, 2.1-megapixel webcam), securely mounted on a tripod at a fixed distance above the tank. A laminated grid sheet with 1 cm intervals was placed at the bottom of the open central arm to facilitate measurements.

Fish were introduced to the holding space and acclimatized in the dark shelter for 2 minutes before the guillotine door was opened, allowing them to explore the open passage for 3 minutes. Videos were analyzed using BORIS behavioral event-logging software (Friard and Gamba 2016) to measure the latency to emerge from the shelter after the guillotine door was lifted (Figure 1).

### Social Preference Test

We conducted a shoal size preference test using a multi-shoal choice setup to infer preferences in two conditions: (1) two shoaling options (4 versus 2 fish) and (2) three shoaling options (4 versus 2 fish, with a third option of 1 fish). All shoals were presented in rectangular display tanks arranged around a circular arena. The subject fish, placed in the arena, could see the shoals in the display tanks through the transparent walls of the circular arena and express their preference by shoaling within one of four sectors, each facing a corresponding rectangular display tank. In the binary condition, shoals were always presented in randomly chosen display tanks positioned opposite each other. In the ternary condition, a decoy was randomly introduced into one of the two empty display tanks.

The setup comprised a cylindrical acrylic focal tank (24 cm in diameter, 4 mm thick) placed at the center of a larger square glass main tank (54 cm × 54 cm × 31 cm) (see apparatus, Figure 1). Rectangular display tanks, each measuring 27 cm in length, 10 cm in width, and 30 cm in height, were positioned 4 cm from the focal tank and attached to the main tank. The entire setup was filled with water to a height of 15 cm. The main tank was elevated on a raised platform, with its external surfaces covered in opaque paper to eliminate external visual distractions. To prevent interactions between the display fish, the perpendicular faces of the display tanks were lined with laminated black paper.

LED light strips were installed inside each display tank to ensure consistent internal illumination. The setup was surrounded by a 10 ft × 10 ft black photography curtain to block external light, maintaining the display tanks as the only light source. A thin layer of water between the display and focal tanks reduced refraction distortion. The background of each display tank, located behind the display fish, was covered with a green laminated paper sheet, as outlined by Lucon-Xiccato, Dadda, et al. (2017). Recordings were made using a GoPro Hero 8 camera (1080p, 30 fps, linear mode).

The movement of fish within each setup was tracked using DeepLabCut (Nath et al. 2019), a deep learning-based software for precise trajectory analysis. A custom R script was developed to process the trajectory data generated by DeepLabCut and to quantify the time spent in each shoal-facing sector. The relative preference for the larger shoal (4 fish) over the smaller shoal (2 fish), both with and without the presence of a third shoal (1 fish), was calculated for two-choice and three-choice conditions using the following equation:

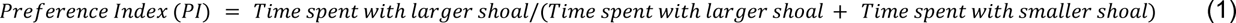

### Experimental Procedure

Each fish underwent a sequence of three personality tests—an Open Field Test (OFT), an Emergence Test (ET), and a Social Preference Test (SPT)—in the same order on any given trial day, with a two-hour time window allocated for each test to control for the time of day. The testing sequence was randomized for each fish within each test. Between tests, the fish were returned to their housing tank near the behavioral testing area to minimize stress during the entire testing period.

This battery of tests was repeated twice over two days, with a two-day gap between repetitions (referred to as Trials), under three different hunger state conditions: H0: Same-day fed (30 minutes before testing), H1: Fed 24 hours before testing, H2: Fed 48 hours before testing. Fish were fed exclusively in the mornings according to these schedules to ensure consistency. Two testing cycles were conducted, separated by a two-week interval, with each cycle consisting of two repetitions of the test battery under each hunger state (Figure 1). This design allowed us to disentangle the effects of hunger state from potential plasticity arising from habituation to the tests and time spent in isolation in holding tanks. Due to time constraints in completing the tests within a consistent time window each day, testing was carried out in four batches of six fish each (24 males).

### Ethical Note

All animal procedures were carried out in compliance with the guidelines set by the Committee for Control and Supervision of Experiments on Animals (CCSEA) and in alignment with national and institutional ethical standards for animal care and use. Ethical clearance was granted by the Institutional Animal Ethics Committee of Ashoka University (approval number ASHOKA/IAEC/2/2022/6).

### Statistical Analysis

We collected a total of 288 behavioral measures for each test across the entire study. The following analysis pipeline was used to address the research questions outlined in the introduction:

1. To evaluate whether individuals differed in their average performance in the behavioral tests, we employed univariate mixed-effects models. In addition to fixed effects, these models included fish ID as a random factor. This approach allowed us to partition the variance attributable to between-individual differences and calculate repeatability for each behavioral test.
2. For behaviors with a significant proportion of variance explained by between-individual differences, we examined whether individuals varied in their responsiveness (plasticity) to changes in hunger state. Slope models were fitted for tests with significant repeatability, enabling us to quantify between individual differences in plasticity for each behavioral test.
3. For behaviors with a significant proportion of variance explained by between-individual differences in the univariate models, we would typically use bivariate models to assess covariances and establish behavioral syndromes. However, in our study, this was unnecessary as no response variable, other than time spent in the open in the Open Field test, showed significant repeatability.

Using the pipeline described above and based on our experimental design, we fitted univariate linear mixed-effects models for each response variable for each behavioral test. The models included hunger state (H0, H1, H2), cycle (two-week gap between test repetitions; see design, Figure 1), trial (two-day gap between repetitions within each hunger state and cycle; see design, Figure 1), and population cohort as fixed effects, with ID and batch as random effects. We fitted these models using the `lmer` function from the ‘lme4’ package in R. Hunger state was treated as a covariate rather than a factor, as modeling it as a covariate provided a better fit, as determined by a likelihood ratio test.

To evaluate the main effect of introducing a decoy option (decoy effect), we fitted a mixed-effects model combining data from binary and ternary preference tests. This model included decoy condition as a fixed effect, along with hunger state, cycle, cohort, and trial as additional fixed effects.

To estimate the fixed effects, we simulated 2000 values from the posterior distributions of the models using the ‘sim’ function in the ‘arm’ package in R. We then extracted 95% confidence intervals (CI) around the mean to assess parameter uncertainty. Model fit was evaluated by visually inspecting the scaled residuals using the ‘DHARMa’ package in R (Hartig 2018). Overdispersion in the emergence test model residuals was corrected by applying a log transformation to the latency to emerge response variable.

Adjusted repeatability (R) was defined as the proportion of phenotypic variance attributed to between-individual differences after accounting for fixed effects, calculated as the ratio of among-individual variance to total variance (Nakagawa and Schielzeth 2010). Predicted values for each response variable were obtained from the univariate models using the predict_response() function from the ggeffects package in R. These predicted values were assessed for any deviation from a chance-level relative preference of 0.5. To assess between-individual variation in plasticity, we fitted a slope model for hunger state, as cycle and cohort had only two levels. We then compared univariate models with and without random slopes using the anova() function from the ‘stats’ package in R.

**Table 1.**
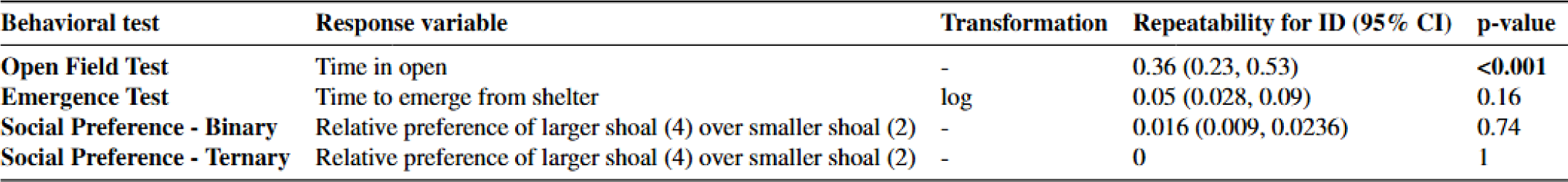
Adjusted repeatability estimates from univariate models.

## Results

### Effect of Hunger state, Cycle, Cohort and Trial on the average behavior

Time spent in the open field was significantly higher in the second cycle compared to the first cycle across both cohorts (*estimate: 11.55, 95% CI: 2.82 to 20.38*). However, the second population cohort spent significantly less time in the open field overall across both cycles (*estimate: −31.94, 95% CI: −54.37 to −7.90*; Figure 2, Table S1). In contrast, hunger state (estimate: 0.124, 95% CI: −5.31, 5.58) and trial number (estimate: −3.44, 95% CI: −12.62, 5.74) did not significantly influence time spent in the open field (Figure 2, Table S1).

**Figure 2.**
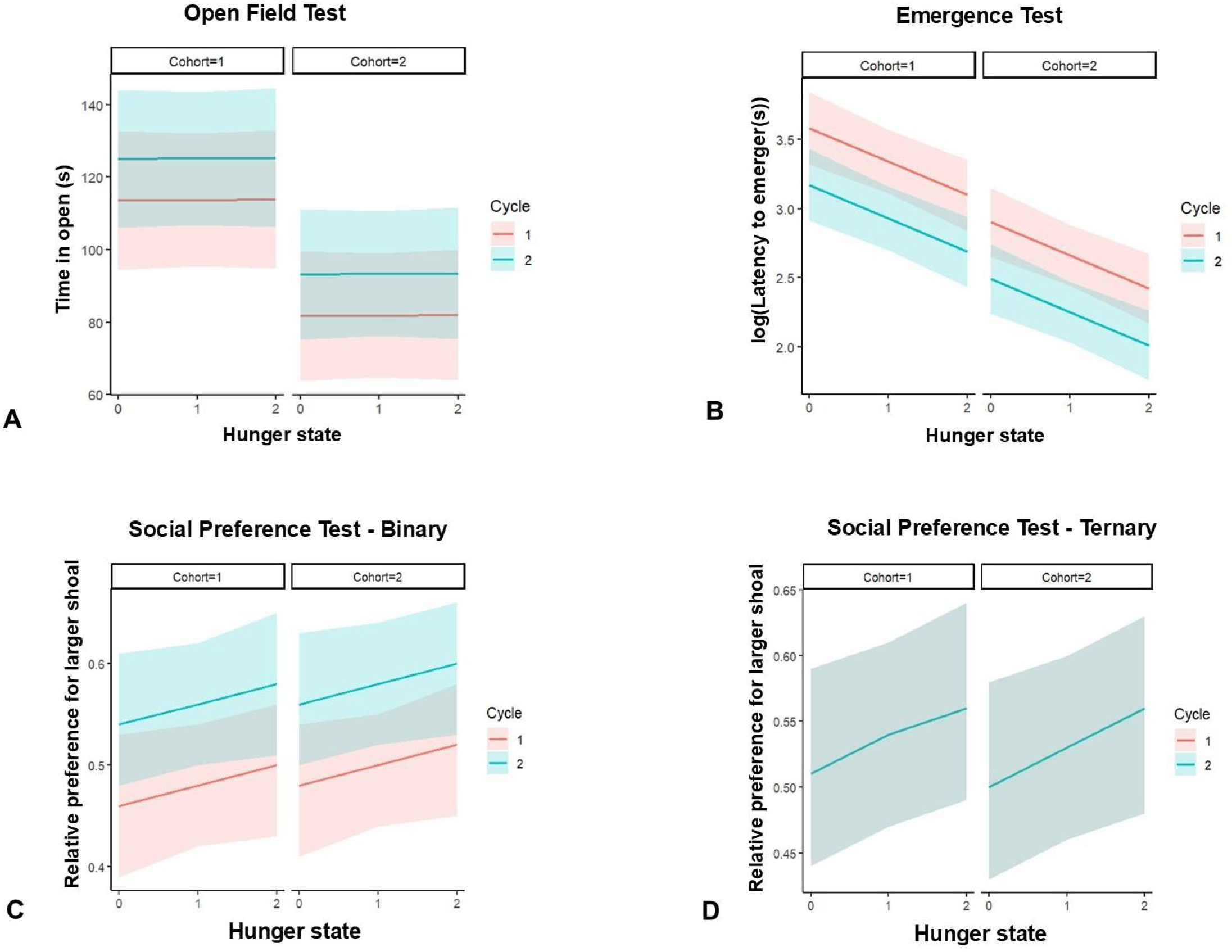
Predicted estimates of fixed effects with 95% confidence intervals (CIs) from univariate models for each response variable across behavioral tests. (A) Time spent in the open (seconds) during the Open Field Test. (B) Log-transformed latency to emerge (seconds) during the Emergence Test. (C) Relative preference for the larger shoal in the Social Preference Test (Binary choice). (D) Relative preference for the larger shoal in the Social Preference Test (Ternary choice).

Log-transformed latency to emerge during the ET decreased significantly with increasing hunger state (*estimate: −0.242, 95% CI: −0.363 to −0.123*), cycle (estimate: −0.408, 95% CI: −0.595, −0.206), and cohort (estimate: −0.677, 95% CI: −0.941, −0.422). Trial number had no significant effect on latency to emerge during the ET (estimate: 0.097, 95% CI: −0.101, 0.287; Figure 2, Table S1).

In the binary shoal choice test, cycle significantly increased the relative preference for the larger shoal (estimate: 0.082, 95% CI: 0.028, 0.135; Figure 2, Table S1). This effect was driven by an increase in time spent with the larger shoal (estimate: 10.39, 95% CI: 3.038, 17.54) and a corresponding decrease in time spent with the smaller shoal (estimate: −13.57, 95% CI: −21.41, −5.77; Table S2). Hunger state (estimate: 0.0182, 95% CI: −0.0150, 0.0507), cohort (estimate: 0.0190, 95% CI: −0.0401, 0.0805), and trial (estimate: −0.0222, 95% CI: −0.0798, 0.0312) did not significantly influence relative preference (Figure 2, Table S1). The predicted relative preference for the larger shoal increased significantly above chance (0.5) during Cycle 2 for both cohorts (Cohort 1: estimate: 0.56, 95% CI: 0.51, 0.62; Cohort 2: estimate: 0.58, 95% CI: 0.52, 0.64) and under both hunger conditions (H1: Cohort 1: estimate: 0.58, 95% CI: 0.51, 0.65; Cohort 2: estimate: 0.60, 95% CI: 0.53, 0.66; Table S3).

In the ternary shoal choice test, none of the predictors significantly influenced the relative preference for the larger shoal, including hunger state (estimate: 0.0263, 95% CI: −0.0032 to 0.0571), cycle (estimate: 0.0001, 95% CI: −0.0529 to 0.0530), cohort (estimate: −0.0087, 95% CI: −0.0982 to 0.0842), or trial (estimate: −0.009, 95% CI: −0.0606 to 0.0410; Figure 2, Table S1). Similarly, none of these predictors significantly influenced the time spent with either the larger or smaller shoal. However, hunger state exhibited a marginal effect on the time spent with the decoy shoal (estimate: 2.84, 95% CI: −0.63 to 6.13), which was accompanied by a marginal decrease in time spent with the smaller shoal (estimate: −3.10, 95% CI: −7.22 to 1.04), with no effect observed for the larger shoal (estimate: 0.94, 95% CI: −3.20 to 4.90; Table S2).

### Individual repeatabilities for each behavior

Significant repeatability was observed only in the open field test, with time in the open field yielding a repeatability estimate of 0.36 (0.23, 0.53). In contrast, all other measured behaviors exhibited near-zero repeatability.

### Individual differences in plasticity

Adding random slopes for hunger state (χ²(2) = 0.68, p = 0.70) did not significantly improve the model fit compared to the individual identity (ID) model for time spent in the open during the Open Field Test (OFT), indicating no significant individual variation in responsiveness to hunger.

### Decoy effect

In the combined model analyzing relative preferences for the larger shoal size (4 over 2), with the decoy option (1 fish) included as a fixed effect, the decoy option had no significant effect (*χ*^2^(1) = 0.62, *p* = 0.43).

## Discussion

### Boldness tests

Hunger state significantly influenced emergence time, with shorter latencies to emerge when hungry, while exploration behavior in the open field test remained unaffected. Cycle, i.e., the two-week interval between tests, increased boldness in both measures, leading to more time spent in the open and shorter latency to emerge. This increased boldness might reflect habituation to the setup or reduced anxiety. The population cohort influenced both boldness measures—time spent in the open and latency to emerge from shelter—but not in a correlated manner. The second cohort spent significantly less time in the open (indicating shyness) but had a lower latency to emerge from shelter (indicating boldness). Among the two boldness tests, only time spent in the open during the Open Field test was significantly repeatable, with a sizeable repeatability.

The differences in cohort effects, hunger state influences, and the repeatability of the measures suggest that the emergence and exploration tests assess distinct behavioral traits. Boldness is commonly assessed using the emergence test, where an animal voluntarily leaves a shelter to explore a novel environment, and the Open Field test, which involves forced exploration of a novel setting or object (Alecia J. Carter et al. 2013). While these tests are often used interchangeably under the assumption that they measure similar or related traits, evidence suggests they may not always correlate and could reflect distinct traits, such as exploration/curiosity versus anxiety (Misslin and Cigrang 1986; Walsh and Cummins 1976; Alecia J. Carter et al. 2013; Yuen et al. 2017; Andersson, Laikre, and Bergvall 2014; Perals et al. 2017).

The lack of an effect of hunger state on boldness in the Open Field test has been previously reported in zebrafish, and our results align with these findings (Ariyomo and Watt 2015). The Open Field test also demonstrates robust repeatability in zebrafish, consistent with earlier studies (H. R. Thomson et al. 2020; Baker et al. 2018). However, to our knowledge, no studies have examined the effect of hunger state on boldness using latency to emerge from a refuge in zebrafish. Evidence from other species suggests that hunger enhances risk-taking behavior (Näslund 2021; J. S. Thomson et al. 2012; Moran et al. 2021). While several studies in zebrafish have used emergence from shelter to assess boldness (Roy and Bhat 2018; Araujo-Silva et al. 2018; Fu, Zhang, and Fan 2024; Vargas, Mackenzie, and Rey 2018), repeatability is not consistently reported. Moreover, the validity of this approach has been questioned due to issues with replicability (Näslund 2021) and its sensitivity to experimental design (Näslund, Bererhi, and Johnsson 2015). Our findings reveal a lack of repeatability in this test and demonstrate that latency to emerge is influenced by population cohort, habituation due to repeated exposure (cycle), and hunger state, underscoring the context dependency of this measure and its limitations for assessing personality traits.

### Social Preference tests

Cycle had a significant and increasing influence on the overall relative preference for the larger shoal (4 vs. 2). Starting from no baseline preference (chance level: 0.5), a significant preference for the larger shoal emerged in Cycle 2, particularly for hungry fish (H1 and H2). This shift was driven by a significant increase in the time spent with the larger shoal, paired with a corresponding decrease in time spent with the smaller shoal in the binary test. Population cohorts showed no significant effect on social preference in either the binary or ternary tests, indicating the robustness of shoal size preference across populations.

In the ternary test, relative preference remained stable, with no preference for the larger shoal observed across any cycle, cohort, or hunger state. A marginal, non-significant increase in time spent with the decoy shoal (single fish) was observed due to hunger state, paired with a marginal decrease in time spent with the smaller shoal. However, hunger state did not significantly influence the time spent with the larger shoal. No repeatability was observed in the relative preference in either the binary or ternary tests.

To our knowledge, no studies have examined the effect of hunger on shoal size preference in zebrafish. In other fish species, such as Three-spined sticklebacks (*Gasterosteus aculeatus*) (Krause 1993) and Golden Shiners (*Notemigonus crysoleucas*) (Reebs and Saulnier 1997), hunger has been shown to decrease social preference. In our study, we observed a marginal increase in shoal size preference due to hunger, compared to chance (0.5), during the second cycle, alongside a much stronger effect of the cycle itself. This effect could be male-specific (to be tested with females), as males generally exhibit weaker preferences for larger shoals. This pattern was also evident in our study, where there was no baseline preference for the larger shoal. The hunger-induced shift toward preferring larger shoals may offer a foraging advantage, as larger shoals can enhance foraging success (Polyakov et al. 2022).

The increase in preference due to the cycle is unlikely to result from social isolation, as adult zebrafish are known to show reduced shoaling preference after isolation (Müller et al. 2024). In our setup, fish were housed in isolated transparent tanks with a shared water supply, preventing visual and olfactory isolation. The observed effect may reflect either habituation or reduced anxiety in response to repeated exposure to the preference task, and our design could not disentangle these possibilities. We maintained the same order for testing, with the binary choice set tested first followed by the ternary choice set, for all fish to control for social exposure and baseline preference capture, while ensuring a sufficient sample size. As such, our conclusions are limited to a single order condition. Notably, no cycle effect was observed in the ternary shoal preference task, which may be due to prior exposure to shoals (order effect) or the stabilizing influence of the decoy shoal. Experimental dissection of these conditions in future studies could clarify their independent contributions.

No decoy effect was observed, consistent with our previous work (Singh et al. 2022), which also reported no decoy effect under similar testing conditions when the binary choice set always preceded the ternary choice set among males. In the ternary choice set, a marginal increase in time spent with the decoy option was observed as hunger increased, paired with a marginal decrease in time spent with the smaller shoal. This resulted in an increased relative preference for the larger shoal. It would be intriguing to further investigate whether this effect could lead to a significant shift in relative preference if tested with a different order or among females. The influence of internal states like hunger on cognitive biases, such as violations of rationality, is of significant interest, as explored in other species (Schuck-Paim, Pompilio, and Kacelnik 2004; Yamada 2017).

### Conclusion

Physiological state, rather than personality, plays a critical role in shaping key behavioral traits in zebrafish males, such as emergence from shelter and shoal size discrimination. The Open Field Test (OFT) proved to be a more reliable assay for assessing boldness as a personality trait, with repeatability observed in the OFT but not in the emergence test. Importantly, no association was found between personality and state, even in the OFT, where hunger had no measurable effect on boldness. The results contribute to a growing body of evidence suggesting limited or no association between state and personality (MacGregor, Cottage, and Ioannou 2021; Parthasarathy et al. 2022; Niemelä and Dingemanse 2018).

This study also provides the first empirical evidence of low repeatability in shoal size discrimination in any fish species. This behavior was found to be influenced by physiological factors, including hunger and repeated task exposure. These findings challenge the generalization of personality traits, from sociability (Gartland et al. 2022), to cognitive measures like shoal size discrimination, cautioning against assuming a common underlying trait. This interpretation aligns with recent studies that reported weak or no correlations between sociability and shoaling preference (Lucon-Xiccato and Dadda 2017; Fu, Zhang, and Fan 2024).

To further disentangle the effects of labile physiological states, such as hunger or anxiety, from personality-driven behaviors, future studies should emphasize experimental designs that actively manipulate these states. This work provides a nuanced understanding of the complexity of behavioral traits in zebrafish, offering valuable insights into behavioral ecology and cognitive research across species.

## Competing Interest

The authors declare no competing interest.

## Acknowledgements

This research was supported by the Ashoka University Annual Faculty Research Grant awarded to BKR, as well as the infrastructure provided by the Research Office at Ashoka University. AS extends gratitude to Jonathan Wright for facilitating the SQUiD fellowship, enabling a visit to NJD’s lab at LMU Munich. Special thanks are also extended to Barbara Class and Irene Goana for their valuable discussions and insights on statistical analysis. We also thank Corné de Groot and Shivani Krishna for reviewing the manuscript.

## Author contributions statement

**Abhishek Singh**: Conceptualization, Methodology, Formal analysis, Investigation, Visualization, Writing Original draft preparation. **Prabhat Sharma**: Methodology, Investigation. **Mansha Jain**: Investigation, Validation. **Pranati Mahajan**: Investigation, Validation. **Jasmine Bajaj**: Investigation, Validation. **Vaidehi Gupta**: Validation. **Niels J. Dingemanse**: Formal analysis. **Bittu K. Rajaraman**: Supervision, Funding acquisition, Writing- Reviewing and Editing.

## Notes

### Competing Interest Statement

The authors have declared no competing interest.

