## Supplementary Information for "State or personality trait: Determinants of boldness and shoal size preference in adult zebrafish"

**Table S1: Model estimates, 95% confidence intervals, and likelihood ratio test results for the fixed effects of each behavioral test.**

**Time spent in open (s) in Open Field Test**

| Effect | Estimate | 95% CI | $\chi^2$ (Chi-square) | P-value |
|---|---|---|---|---|
| Hunger state | 0.12 | (-5.31, 5.58) | 0.002 | 0.95 |
| Cycle | **11.55** | **(2.82, 20.38)** | **6.45** | **0.01** |
| Cohort | **-31.94** | **(-54.37, -7.90)** | **7.15** | **0.007** |
| Trial | -3.44 | (-12.62, 5.74) | 0.59 | 0.43 |

**Log latency to emerge from shelter (s) in Emergence Test**

| Effect | Estimate | 95% CI | $\chi^2$ (Chi-square) | P-value |
|---|---|---|---|---|
| Hunger state | **-0.242** | **(-0.363, -0.123)** | **16.22** | **<0.001** |
| Cycle | **-0.408** | **(-0.595, -0.206)** | **17.01** | **<0.001** |
| Cohort | **-0.677** | **(-0.941, -0.422)** | **28.18** | **<0.001** |
| Trial | 0.097 | (-0.10, 0.287) | 0.94 | 0.332 |

**Relative preference for the larger shoal (4) over smaller shoal (2)- binary test**

| Effect | Estimate | 95% CI | $\chi^2$ (Chi-square) | P-value |
|---|---|---|---|---|
| Hunger state | 0.0182 | (-0.0150, 0.0507) | 1.22 | 0.269 |
| Cycle | **0.0820** | **(0.0283, 0.1355)** | **8.79** | **0.003** |
| Cohort | 0.0190 | (-0.0401, 0.0805) | 0.37 | 0.546 |
| Trial | -0.0222 | (-0.0798, 0.0312) | 0.65 | 0.41 |

**Relative preference for the larger shoal (4) over smaller shoal (2)- ternary test**

| Effect | Estimate | 95% CI | χ² (Chi-square) | P-value |
|---|---|---|---|---|
| Hunger state | 0.026 | (-0.0032, 0.057) | 2.59 | 0.10 |
| Cycle | 0.0001 | (-0.053, 0.053) | <0.01 | 0.98 |
| Cohort | -0.0087 | (-0.098, 0.084) | 0.04 | 0.85 |
| Trial | -0.009 | (-0.06, 0.04) | 0.11 | 0.74 |

**Table S2: Model estimates, 95% confidence intervals, and likelihood ratio test results for the fixed effects on time spent in zones during the social preference tests.**

**Time spent with larger shoal (4) in binary test**

| Effect | Estimate | 95% CI | χ² (Chi-square) | P-value |
|---|---|---|---|---|
| Hunger state | 2.87 | (-1.67, 7.32) | 1.54 | 0.213 |
| Cycle | **10.39** | **(3.04, 17.54)** | **7.19** | **0.007** |
| Cohort | 7.82 | (-2.22, 17.11) | 2.43 | 0.118 |
| Trial | -3.27 | (-10.43, 3.90) | 0.74 | 0.388 |

**Time spent with larger shoal (2) in binary test**

| Effect | Estimate | 95% CI | χ² (Chi-square) | P-value |
|---|---|---|---|---|
| Hunger state | -2.79 | (-7.6,1.64) | 1.40 | 0.23 |
| Cycle | **-13.57** | **(-21.40,-5.77)** | **11.53** | **0.0006** |
| Cohort | 3.42 | (-5.31,12.12) | 0.56 | 0.45 |
| Trial | 2.65 | (-4.98,10.20) | 0.46 | 0.50 |

**Time spent with larger shoal (4) in ternary test**

| Effect | Estimate | 95% CI | $\chi^2$ (Chi-square) | P-value |
|---|---|---|---|---|
| Hunger state | 0.9433 | (-3.201,4.895) | 0.23 | 0.62 |
| Cycle | -2.0495 | (-8.837,4.852) | 0.31 | 0.57 |
| Cohort | -0.0624 | (-12.737,13.481) | 0.0004 | 0.98 |
| Trial | -0.0906 | (-7.032,6.826) | 0.0001 | 0.99 |

**Time spent with smaller shoal (2) in ternary test**

| Effect | Estimate | 95% CI | $\chi^2$ (Chi-square) | P-value |
|---|---|---|---|---|
| Hunger state | -3.09 | (-7.22,1.0) | 2.34 | 0.12 |
| Cycle | -0.60 | (-7.01,6.32) | 0.046 | 0.82 |
| Cohort | -0.90 | (-7.64,5.85) | 0.05 | 0.81 |
| Trial | 1.69 | (-4.67,8.59) | 0.24 | 0.62 |

**Time spent with decoy shoal (1) in ternary test**

| Effect | Estimate | 95% CI | $\chi^2$ (Chi-square) | P-value |
|---|---|---|---|---|
| Hunger state | 2.8404 | (-0.6292,6.1314) | 2.80 | 0.009 |
| Cycle | 0.00317 | (-5.4025,5.3696) | 0.0005 | 0.98 |
| Cohort | 4.1346 | (-1.7268,9.3910) | 2.17 | 0.14 |
| Trial | 1.0726 | (-4.3391,6.4939) | 0.14 | 0.70 |

**Table S3: Predicted values of relative preference for the larger shoal from the univariate model of SPT binary across cycles, cohorts, and hunger states**

| Cycle | Cohort | Hunger state | Predicted value | 95% CI |
|---|---|---|---|---|
| 1 | 1 | 0 | 0.46 | 0.39, 0.53 |
| 1 | 1 | 1 | 0.48 | 0.42, 0.54 |
| 1 | 1 | 2 | 0.50 | 0.43, 0.56 |
| 1 | 2 | 0 | 0.48 | 0.41, 0.54 |
| 1 | 2 | 1 | 0.50 | 0.44, 0.55 |
| 1 | 2 | 2 | 0.52 | 0.45, 0.58 |
| 2 | 1 | 0 | 0.54 | 0.48, 0.61 |
| 2 | 1 | 1 | **0.56** | **0.51, 0.62** |
| 2 | 1 | 2 | **0.58** | **0.51, 0.65** |
| 2 | 2 | 0 | 0.56 | 0.50, 0.63 |
| 2 | 2 | 1 | **0.58** | **0.52, 0.64** |
| 2 | 2 | 2 | **0.60** | **0.53, 0.66** |

Values in bold indicate a significantly above-chance preference (0.5) for the larger shoal (4 vs. 2); Values adjusted for Trial = 1, ID = 0, and Batch = 0.

**Table S4: Predicted values of relative preference for the larger shoal from the univariate model of SPT ternary across cycles, cohorts, and hunger states**

| Cycle | Cohort | Hunger state | Predicted value | 95% CI |
|---|---|---|---|---|
| 1 | 1 | 0 | 0.51 | 0.44, 0.59 |
| 1 | 1 | 1 | 0.54 | 0.47, 0.61 |
| 1 | 1 | 2 | 0.56 | 0.49, 0.64 |
| 1 | 2 | 0 | 0.50 | 0.43, 0.58 |
| 1 | 2 | 1 | 0.53 | 0.46, 0.60 |
| 1 | 2 | 2 | 0.56 | 0.48, 0.63 |
| 2 | 1 | 0 | 0.51 | 0.44, 0.59 |
| 2 | 1 | 1 | 0.54 | 0.47, 0.61 |
| 2 | 1 | 2 | 0.56 | 0.49, 0.64 |
| 2 | 2 | 0 | 0.51 | 0.43, 0.58 |
| 2 | 2 | 1 | 0.53 | 0.46, 0.60 |
| 2 | 2 | 2 | 0.56 | 0.48, 0.63 |

Values adjusted for Trial = 1, ID = 0, and Batch = 0.
